# Do nocturnal pollinators carry a more conspecific pollen load than diurnal ones?

**DOI:** 10.1101/2023.11.30.569345

**Authors:** Mialy Razanajatovo, Frank M. Schurr, Nadia Muhthassim, Sandra Troesch, Eva Knop

**Affiliations:** Institute of Landscape and Plant Ecology, University of Hohenheim, Ottilie-Zeller-Weg 2, D-70599, Stuttgart, Germany; KomBioTa - Center for Biodiversity and Integrative Taxonomy, University of Hohenheim & State Museum of Natural History, Wollgrasweg 23, D-70599, Stuttgart, Germany; Pädagogische Hochschule Bern, Fabrikstrasse 8, 3012 Bern, Switzerland; ETOP IBCSG Partners Foundation, Effingerstrasse 33, 3008 Bern, Switzerland; Agroscope, Reckenholzstrasse 191, 8904 Zürich, Switzerland; Department of Evolutionary Biology and Environmental Studies, University of Zurich, Winterthurerstrasse 190, 8057 Zürich, Switzerland

**Keywords:** Biotic interactions, circadian cycles, community ecology, ecosystem services, floral ecology, pollination ecology

## Abstract

Plant-pollinator interactions are key for the reproduction of wild plants and for food security. However, the role nocturnal pollinators play in wild plant communities is not yet clear. Specifically, it has rarely been studied whether nocturnal pollinators are comparable to diurnal ones in the pollination services they deliver in plant communities. We tested whether nocturnal pollinators have the potential to provide higher pollination services than diurnal ones by carrying a more conspecific pollen load. We compared pollen loads carried by nocturnal and diurnal pollinators captured over 24-hour cycles in co-flowering plant communities in European ruderal meadows. Pollen load was less diverse at night, and the proportion of conspecific pollen carried by nocturnal pollinators was similar to or higher than that of diurnal ones. Thus, nocturnal pollinators do not only carry pollen but can remove and deposit conspecific pollen with a comparable or even superior performance to diurnal ones. Therefore, nocturnal pollinators can be more efficient pollination vectors than diurnal ones, which might compensate their lower pollen load.

## Introduction

Plant-pollinator interactions are key mutualisms for the reproduction of wild plants and for food security. About half of all flowering plant species depend on pollinators for more than 80% of their seed production (Rodger et al., 2021). Furthermore, more than 80% of crops including vegetables, fruits and stimulants depend on pollinators (Klein et al., 2007). However, declines in pollinating insects associated with global change drivers such as climate change, land-use change, agricultural intensification, invasive species, and spread of pathogens are widespread (Potts et al., 2010). Recent evidence suggests that increasing light pollution which disproportionally affects nocturnal insects adds to the drivers of pollinator declines and their pollination service (Giavi et al., 2021; Knop et al., 2017). As pollinators face the threats of many global change drivers, declines in pollinator abundance and diversity can have cascading impacts on plant communities (Clough et al., 2014). Since studies on plant-pollinator interactions are largely biased towards diurnal interactions, we specifically lack knowledge of nocturnal interactions in a community context (Knop et al., 2018, 2017; Macgregor et al., 2017). In particular, it is unclear which roles nocturnal pollinators play for wild plant communities and for cropping systems (Buxton et al., 2022; Hahn and Brühl, 2016).

Previous studies on nocturnal interactions were limited to very specialized interactions or involving specific groups of pollinators or plants, and most studies were done in tropical regions. The importance of nocturnal pollination by insects in temperate regions has only been recognized recently. A few previous studies in temperate regions focus on moths and suggest their potential roles in plant-pollinator networks by showing that moths captured in light traps carry pollen.Three to 10 %, 76% and 20.7% of captured moths in a boreal forest in Scotland (Devoto et al., 2011), in a biodiversity hotspot in Portugal (Banza et al., 2015) and in the Balearic Islands (Ribas-Marquès et al., 2022), respectively, carried pollen. In the Himalayas, 65% of settling moths from which pollen was extracted from proboscices were considered potential pollinators (Singh et al., 2022). To promote plant reproduction by outcrossing, moths must not only carry pollen but also transfer it to conspecific plants. As heterospecific pollen transfer can interfere with the fertilization of ovules by conspecific pollen by mechanical clogging and allelopathy, and can reduce seed production (Ashman and Arceo-Gómez, 2013), a higher proportion of conspecific pollen transfer can be more beneficial for plants. In an experiment using fluorescent pollen-tracker powder, Buxton et al. (2021) showed that moths can transfer pollen among flowers of the same species. But we still lack knowledge of conspecific pollen transfer by nocturnal pollinators at the community level.

Due to the difference in abiotic conditions (e.g. cooler temperature and darker conditions at night), pollinator communities and therefore plant-pollinator interactions differ notably between day and night. In Swiss ruderal meadows, Knop et al. (2018) found that the dominant groups of pollinators at night (defined based on whether they carried pollen or not) were Lepidopterans and Coleopterans, whereas diurnal pollinators were mostly Dipterans and Hymenopterans. Also, fewer pollinator species and individuals visited flowers at night than during the day: 16% of pollinator species visited flowers only at night, and 9% did so during both night and day (Knop et al., 2018). However, whether nocturnal pollinators interact with a more or less diverse set of plant species than diurnal ones has not been studied yet.

Pollen carried externally by pollinators can reflect the assemblage of plant species that they visit (Kearns and Inouye, 1993). Moreover, conspecific pollen removal by the pollinator as well as the potential for conspecific pollen deposition can be derived from the proportion of pollen grains from the visited flower that are observed on the pollinator body. To our knowledge, no study compared pollen loads carried by nocturnal and diurnal pollinators in co-flowering plant communities and tested whether nocturnal pollinators carried more conspecific pollen than diurnal ones. Here, we compared pollen loads carried on the body of nocturnal and diurnal flower-visiting insects in Swiss ruderal meadows during 24-hour cycles. Specifically, we asked whether the pollen load, the number of pollen species, the amount and proportion of conspecific pollen carried by pollinators vary across a 24-hour cycle.

## Methods

### Pollen load dataset

We used a dataset of pollen load carried by flower visitors captured over 24-hour cycles in European ruderal meadows located in eight sites between 709 and 851 m a.s.l. (Knop et al., 2018, 2017). Between June and August 2014, we observed and captured insects actively visiting open flowers along one 50m transect per site, with up to six repetitions per transect, on days and nights without rain and strong wind. We only considered insects touching the reproductive parts of the plant as flower visitors (i.e., excluded the ones destroying the petals or sleeping in the flower). We removed the pollen from the insect bodies, and identified pollen species under light microscope using our own pollen reference database. We built the pollen reference database using plant species found around the transect. We collected pollen load data from 3,626 insects belonging to 500 species from 98 families and 10 orders, caught visiting flowers of 99 plant species. In total, we identified pollen grains of 127 plant species on the flower visitors.

### Data analysis

To differentiate potential pollinators from mere flower visitors, we classified all insect individuals that carried five or more pollen grains of any plant species as potential pollinators (Devoto et al., 2011; Knop et al., 2017). To analyze the variation in pollen load over 24-hour cycles, we ran a generalized linear mixed model with a negative binomial distribution using the glmer.nb function of the lme4 package (Bates et al., 2014) in R version 4.3.0 (R-Core-Team, 2023). As response variable, we included the number of pollen grains carried by the pollinators. To account for the cyclical nature of time of day or night, we used the sine and cosine of both *t* and 2\**t* as explanatory variables, where *t* is time in units of radians (Knop et al., 2018). To account for potential variations due to site characteristics and abiotic conditions, we included site and date of capture as random variables.

To compare the diversity of pollen species carried by nocturnal and diurnal pollinators, we ran a generalized linear mixed model with a Poisson distribution using the glmer function of the lme4 package. As response variable, we included the number of pollen species carried by pollinators. As in the previous model, we included the sine and cosine of time of day or night in radians as explanatory variables. To account for potential bias due to variation in pollen quantity carried by different pollinators, we included the log-tranformed total number of pollen grains on each pollinator as a covariate. We also included site and date as random variables.

To analyze the variation in the amount of conspecific pollen carried by pollinators (i.e. pollen from the same plant species they visited), we ran a generalized linear mixed model with a negative binomial distribution using the glmer.nb function of the lme4 package. As response variable, we included the number of conspecific pollen grains carried by the pollinators. As in the previous models, we included the sine and cosine of time of day or night in radians as explanatory variables. To account for potential variations due to site characteristics, we included site as random variable. To test whether nocturnal pollinators carried a higher proportion of conspecific pollen than diurnal ones, we ran a generalized linear mixed model with a binomial distribution using the glmer function of the lme4 package. As response variable, we included the proportion of conspecific pollen grains carried by pollinators. As in the previous models, we included the sine and cosine of time of day or night in radians as explanatory variables, and site as a random variable. Again, we considered only pollinators (carrying > 5 pollen grains of any given plant species) in this analysis. To check the fit of all models, we used the simulateResiduals function in the DHARMa package (Hartig, 2022). For all models, we reported the marginal and conditional *r*^*2*^ (Nakagawa et al., 2017).

## Results

We identified 1,322,737 pollen grains carried by 3,626 flower visitors of which 14.34% were captured at night, i.e. between 22:00 and 05:59. Each flower visitor carried between 0 and 20,742 pollen grains from up to 18 plant species. Pollen load, i.e. the number of pollen grains carried by pollinators, significantly depended on time of day or night over a 24-hour cycle (Table 1). Pollinators carried fewer pollen grains at night (Fig. 1A).

**Table 1.** Results of four generalized linear mixed models testing how the number of pollen grains, the number of pollen species, the amount of conspecific pollen, and the proportion of conspecific pollen carried by pollinators depend on time of day or night (*t* in radians). Significant model parameters are highlighted in bold (p<0.05).

| Response variables | Number of pollen grains | Number of pollen species | Amount of conspecific pollen | Proportion of conspecific pollen |
| --- | --- | --- | --- | --- |
| Fixed terms | Estimate (standard error) |  |  |  |
| Intercept | **5.162 (0.244)** | **1.497 (0.059)** | **4.647 (0.106)** | 0.067 (0.113) |
| $\sin(t)$ | **0.777 (0.057)** | **0.105 (0.023)** | **0.541 (0.096)** | **-0.112 (0.007)** |
| $\cos(t)$ | **-0.192 (0.074)** | **-0.117 (0.029)** | **-0.481 (0.144)** | **-0.012 (0.010)** |
| $\sin(2*t)$ | **0.043 (0.053)** | **0.045 (0.020)** | **-0.458 (0.097)** | **-0.081 (0.006)** |
| $\cos(2*t)$ | **0.088 (0.049)** | 0.022 (0.017) | 0.001 (0.091) | **0.127 (0.005)** |
| Total number of pollen grains |  | **0.240 (0.009)** |  |  |
| Random terms | Standard deviation |  |  |  |
| Site | 0.418 | 0.136 | 0.159 | 0.321 |
| Date | 0.861 | 0.123 |  |  |
| Residuals | 1.000 | 0.929 | 0.870 | 19.155 |
| Marginal $r^2$ | 0.093 | 0.240 | 0.183 | 0.111 |
| Conditional $r^2$ | 0.547 | 0.351 | 0.194 | 0.944 |

**Fig. 1.**
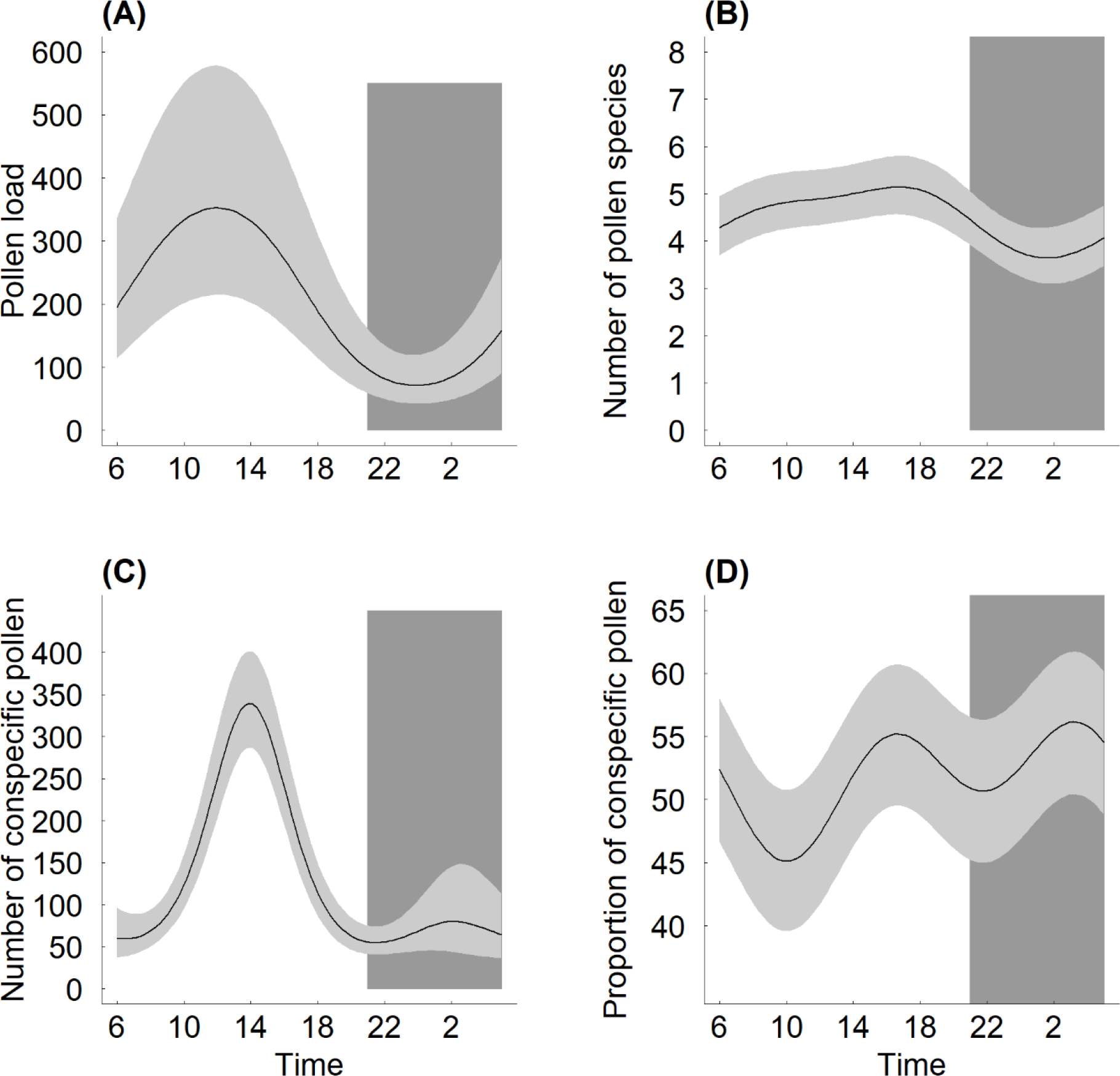
Time of day or night affects (A) pollen load (the number of pollen grains carried by pollinators), (B) the number of pollen species, (C) the amount of conspecific pollen, and (D) and the proportion of conspecific pollen carried by pollinators in Swiss ruderal meadows. Black curves are predictions of generalized linear mixed models and grey areas are 95% confidence intervals.

Time of day or night significantly affected variation in the number of pollen species carried by pollinators (Table 1). Over a 24-hour cycle, the number of pollen species carried by pollinators slowly increased from 6:00 and peaked in the afternoon, then decreased to reach minimal values at night (Fig. 1B). Overall, 72 % of the pollinators carried conspecific pollen, i.e. pollen from the same plant species they visited and on which they were caught. Time of day or night significantly affected variation in amount and proportion of conspecific pollen (Table 1). Over a 24-hour cycle, proportion of conspecific pollen was always larger than 40%, and peaked twice, once in the afternoon and once at night (Fig. 1D).

## Discussion

Our analysis of pollen load carried by pollinators showed that pollen load fluctuated substantially across a 24-hour cycle, with fewer and less diverse pollen carried at night. A lower pollen diversity at night suggests that nocturnal pollinators have a lower linkage density in plant-pollinator networks and play therefore a different role from diurnal pollinators. Also, proportion of conspecific pollen carried by nocturnal pollinators was similar to or higher than that of diurnal ones. This suggests that nocturnal pollinators can be more efficient than diurnal ones, which might partially compensate their lower pollen load.

### Availability of floral resources during a 24-hour cycle

The finding that fewer pollen were collected at night is in accordance with Borges et al. (2016), who suggest that the majority of nocturnal pollinators collect nectar, whereas only fewer ones collect pollen. A smaller and less diverse pollen load at night could be explained by the variation of floral resources available to flower visiting insects over 24 hours. As most temperate plants open new flowers during the day, available floral resources at night are mainly nectar and pollen of flowers which have been already exposed to diurnal pollinators (Willmer, 2011). While plants can adjust nectar production in response to consumption (Heil, 2011), pollen is not replenished (Hargreaves et al., 2009). Furthermore, many plant species close their flowers at night (van Doorn and van Meeteren, 2003), limiting the access to floral resources for nocturnal insects, which might be due to temporal niche partitioning among pollinators, and an avoidance of heterospecific pollen transfer among plants (Fründ et al., 2011). Future studies should explicitly compare the availability of floral resources between day and night.

### The role of insect and plant traits

The finding that nocturnal pollinators carried a less diverse pollen load, even after accounting for variations due to pollen abundance (i.e. smaller pollen load at night), may be explained by variations in insect size and insect-plant trait matching. Generally, larger insects are considered to carry more pollen (e.g. (Földesi et al., 2021)). However, as we do not have data on body mass, we cannot test this. Also, as moths and beetles were the dominant pollinator groups at night, we do not believe that this was the main explanation as both groups contain many large-sized species. Moths that visit bee flowers at night did not touch the anthers with body parts other than their tongues, and therefore may not have removed pollen as it was suggested by Willmer (2011). For example, the Geometridae moth *Perizoma alchemillata* was caught visiting the flower of *Stachys sylvatica* (Lamiaceae) at night, but did not carry the pollen of this plant species. On the other hand, during the day *Bombus pascuorum* (Apidae) visited and carried the pollen of *S. sylvatica*, as was expected for Lamiaceae of which flowers usually present a bilabiate floral structure adapted to pollination by bees (Claßen-Bockhoff, 2007). Because the lack of mechanical fit between the nocturnal insects and flowers of diurnally visited flowers might limit their ability to remove or deposit pollen, it could be that only pollen grains from perfectly matching plant species were found on nocturnal pollinators.

### Nocturnal pollinator performance

Proportion of conspecific pollen in the pollen load can provide information on pollinator performance, as it should positively affect plant reproduction and fitness. The pattern of variation in proportion of conspecific pollen over a 24-hour cycle in our study can be attributed to the pollinator communities. The moderate proportion of conspecific pollen in the morning corresponded to the peak of Diptera visits, and the highest proportion of conspecific pollen in the afternoon and at night corresponded to peaks of Hymenoptera and Lepidoptera visits, respectively (Knop et al., 2018). Even though many Dipterans can have high pollination performance (Doyle et al., 2020; Raguso, 2020), nocturnal pollinators may have on average higher performance, which is comparable to that of Hymenoptera. Thus, nocturnal pollinators can remove or deposit conspecific pollen at least as well as diurnal pollinators do.

### Complementarity between diurnal and nocturnal pollinators

Nocturnal pollinators may play a complementary role to diurnal ones to achieve higher pollination output in plants. Such a complementarity (Blüthgen and Klein, 2011) among diurnal and nocturnal pollinators has been shown for single plant species. In the arid regions of Mexico, to alleviate competition for bat visitation with other coflowering species at night, the cactus *Marginatocereus marginatus* is also pollinated by hummingbirds during the day (Dar et al., 2006). In the common milkweed *Asclepias syriaca* in the temperate meadows of North America, low-frequency high-quality pollination (with outcross-pollen) by moths at night is complemented by low-quality pollination by bumblebees during the day, which have more frequent visitation and remove more pollinia, but transfer more self-pollen (Jennersten and Morse, 1991). In the tropical cloud forests of Ecuador, high-frequency bat pollination at night frequently accompanied with heterospecific pollen transfer in *Aphelandra acanthus* is complemented by pollination by hummingbirds during the day (Muchhala et al., 2009). In the Swiss pre-alps, Knop et al. (2017) found that a reduction of nocturnal pollinators of *Cirsium oleraceum* due to artificial light at night could not be compensated by the diurnal pollinators. Similarly, in the Swiss alps, Alison et al. (2022) found that the clover *Trifolium pratense* is visited by bumblebees during the day and by moths at night, and both moth and bumblebee visitation increased seed set. However, more studies are needed to find out to what extent nocturnal and diurnal pollinators are indeed complementary in a community context.

## Conclusions

Nocturnal pollinators do not only carry pollen but can remove and deposit conspecific pollen with a comparable or even superior performance to diurnal ones. Plant communities may rely on the functional and environmental complementarity between diurnal and nocturnal pollinators for pollination assurance. Given the worldwide pollinator decline and specific risks to nocturnal pollinators (such as light pollution), there is thus an urgent need for studies clarifying the contribution of nocturnal and diurnal pollinators to plant reproduction.

## References

Alison, J., Alexander, J.M., Diaz Zeugin, N., Dupont, Y.L., Iseli, E., Mann, H.M.R., Høye, T.T., 2022. Moths complement bumblebee pollination of red clover: a case for day-and-night insect surveillance. Biology Letters 18, 20220187. 10.1098/rsbl.2022.0187

Ashman, T.-L., Arceo-Gómez, G., 2013. Toward a predictive understanding of the fitness costs of heterospecific pollen receipt and its importance in co-flowering communities. American Journal of Botany 100, 1061–1070. 10.3732/ajb.1200496

Banza, P., Belo, A.D.F., Evans, D.M., 2015. The structure and robustness of nocturnal Lepidopteran pollen-transfer networks in a Biodiversity Hotspot. Insect Conservation and Diversity 8, 538–546. 10.1111/icad.12134

Bates, D., Mächler, M., Bolker, B., Walker, S., 2014. Fitting linear mixed-effects models using lme4. arXiv preprint arXiv:1406.5823 67, 1–48.

Blüthgen, N., Klein, A.-M., 2011. Functional complementarity and specialisation: The role of biodiversity in plant–pollinator interactions. Basic and Applied Ecology 12, 282–291. 10.1016/j.baae.2010.11.001

Borges, R.M., Somanathan, H., Kelber, A., 2016. Patterns and Processes in Nocturnal and Crepuscular Pollination Services. The Quarterly Review of Biology 91, 389–418. 10.1086/689481

Buxton, M., Anderson, B., Lord, J., 2021. Moths can transfer pollen between flowers under experimental conditions [WWW Document]. NZES. URL https://newzealandecology.org/nzje/3457 (accessed 4.19.22).

Buxton, M.N., Gaskett, A.C., Lord, J.M., Pattemore, D.E., 2022. A global review demonstrating the importance of nocturnal pollinators for crop plants. Journal of Applied Ecology 59, 2890–2901. 10.1111/1365-2664.14284

Claßen-Bockhoff, R., 2007. Floral Construction and Pollination Biology in the Lamiaceae. Annals of Botany 100, 359–360. 10.1093/aob/mcm157

Clough, Y., Ekroos, J., Báldi, A., Batáry, P., Bommarco, R., Gross, N., Holzschuh, A., Hopfenmüller, S., Knop, E., Kuussaari, M., Lindborg, R., Marini, L., Öckinger, E., Potts, S.G., Pöyry, J., Roberts, S.P., Steffan-Dewenter, I., Smith, H.G., 2014. Density of insect-pollinated grassland plants decreases with increasing surrounding land-use intensity. Ecology Letters 17, 1168–1177. 10.1111/ele.12325

Dar, S., Arizmendi, Ma. del C., Valiente-Banuet, A., 2006. Diurnal and Nocturnal Pollination of Marginatocereus marginatus (Pachycereeae: Cactaceae) in Central Mexico. Annals of Botany 97, 423–427. 10.1093/aob/mcj045

Devoto, M., Bailey, S., Memmott, J., 2011. The ‘night shift’: nocturnal pollen-transport networks in a boreal pine forest. Ecological Entomology 36, 25–35. 10.1111/j.1365-2311.2010.01247.x

Doyle, T., Hawkes, W.L.S., Massy, R., Powney, G.D., Menz, M.H.M., Wotton, K.R., 2020. Pollination by hoverflies in the Anthropocene. Proceedings of the Royal Society B: Biological Sciences 287, 20200508. 10.1098/rspb.2020.0508

Földesi, R., Howlett, B.G., Grass, I., Batáry, P., 2021. Larger pollinators deposit more pollen on stigmas across multiple plant species—A meta-analysis. Journal of Applied Ecology 58, 699–707. 10.1111/1365-2664.13798

Fründ, J., Dormann, C.F., Tscharntke, T., 2011. Linné’s floral clock is slow without pollinators – flower closure and plant-pollinator interaction webs. Ecology Letters 14, 896–904. 10.1111/j.1461-0248.2011.01654.x

Giavi, S., Fontaine, C., Knop, E., 2021. Impact of artificial light at night on diurnal plant-pollinator interactions. Nat Commun 12, 1690. 10.1038/s41467-021-22011-8

Hahn, M., Brühl, C.A., 2016. The secret pollinators: an overview of moth pollination with a focus on Europe and North America. Arthropod-Plant Interactions 10, 21–28. 10.1007/s11829-016-9414-3

Hargreaves, A.L., Harder, L.D., Johnson, S.D., 2009. Consumptive emasculation: the ecological and evolutionary consequences of pollen theft. Biological Reviews 84, 259–276. 10.1111/j.1469-185X.2008.00074.x

Hartig, F., 2022. DHARMa: Residual Diagnostics for Hierarchical (Multi-Level / Mixed) Regression Models. R package version 0.4.5.

Heil, M., 2011. Nectar: generation, regulation and ecological functions. Trends in Plant Science 16, 191–200. 10.1016/j.tplants.2011.01.003

Jennersten, O., Morse, D.H., 1991. The Quality of Pollination by Diurnal and Nocturnal Insects Visiting Common Milkweed, Asclepias syriaca. The American Midland Naturalist 125, 18–28. 10.2307/2426365

Kearns, C.A., Inouye, D.W., 1993. Techniques for pollination biologists. Techniques for pollination biologists.

Klein, A.-M., Vaissière, B.E., Cane, J.H., Steffan-Dewenter, I., Cunningham, S.A., Kremen, C., Tscharntke, T., 2007. Importance of pollinators in changing landscapes for world crops. Proceedings of the Royal Society B: Biological Sciences 274, 303–313. 10.1098/rspb.2006.3721

Knop, E., Gerpe, C., Ryser, R., Hofmann, F., Menz, M.H.M., Trösch, S., Ursenbacher, S., Zoller, L., Fontaine, C., 2018. Rush hours in flower visitors over a day–night cycle. Insect Conservation and Diversity 11, 267–275. 10.1111/icad.12277

Knop, E., Zoller, L., Ryser, R., Gerpe, C., Hörler, M., Fontaine, C., 2017. Artificial light at night as a new threat to pollination. Nature 548, 206–209. 10.1038/nature23288

Macgregor, C.J., Evans, D.M., Fox, R., Pocock, M.J.O., 2017. The dark side of street lighting: impacts on moths and evidence for the disruption of nocturnal pollen transport. Global Change Biology 23, 697–707. 10.1111/gcb.13371

Muchhala, N., Caiza, A., Vizuete, J.C., Thomson, J.D., 2009. A generalized pollination system in the tropics: bats, birds and Aphelandra acanthus. Annals of Botany 103, 1481–1487. 10.1093/aob/mcn260

Nakagawa, S., Johnson, P.C.D., Schielzeth, H., 2017. The coefficient of determination R2 and intra-class correlation coefficient from generalized linear mixed-effects models revisited and expanded. Journal of The Royal Society Interface 14, 20170213. 10.1098/rsif.2017.0213

Potts, S.G., Biesmeijer, J.C., Kremen, C., Neumann, P., Schweiger, O., Kunin, W.E., 2010. Global pollinator declines: trends, impacts and drivers. Trends in Ecology & Evolution 25, 345–353. 10.1016/j.tree.2010.01.007

Raguso, R.A., 2020. Don’t forget the flies: dipteran diversity and its consequences for floral ecology and evolution. Appl Entomol Zool 55, 1–7. 10.1007/s13355-020-00668-9

R-Core-Team, 2023. A Language and Environment for Statistical Computing. R Foundation for Statistical Computing, Vienna, Austria. <https://www.R-project.org/>.

Ribas-Marquès, E., Díaz-Calafat, J., Boi, M., 2022. The role of adult noctuid moths (Lepidoptera: Noctuidae) and their food plants in a nocturnal pollen-transport network on a Mediterranean island. J Insect Conserv 26, 243–255. 10.1007/s10841-022-00382-7

Rodger, J.G., Bennett, J.M., Razanajatovo, M., Knight, T.M., Kleunen, M. van, Ashman, T.-L., Steets, J.A., Hui, C., Arceo-Gómez, G., Burd, M., Burkle, L.A., Burns, J.H., Durka, W., Freitas, L., Kemp, J.E., Li, J., Pauw, A., Vamosi, J.C., Wolowski, M., Xia, J., Ellis, A.G., 2021. Widespread vulnerability of flowering plant seed production to pollinator declines. Science Advances 7, doi: 10.1126/sciadv.abd3524. https://doi.org/10.1126/sciadv.abd3524

Singh, N., Lenka, R., Chatterjee, P., Mitra, D., 2022. Settling moths are the vital component of pollination in Himalayan ecosystem of North-East India, pollen transfer network approach revealed. Sci Rep 12, 2716. 10.1038/s41598-022-06635-4

van Doorn, W.G., van Meeteren, U., 2003. Flower opening and closure: a review. Journal of Experimental Botany 54, 1801–1812. 10.1093/jxb/erg213

Willmer, P., 2011. Pollination and Floral Ecology, Pollination and Floral Ecology. Princeton University Press. 10.1515/9781400838943

